# Virus genomes reveal the factors that spread and sustained the West African Ebola epidemic

**DOI:** 10.1101/071779

**Authors:** Gytis Dudas, Luiz Max Carvalho, Trevor Bedford, Andrew J. Tatem, Guy Baele, Nuno Faria, Daniel J. Park, Jason Ladner, Armando Arias, Danny Asogun, Filip Bielejec, Sarah Caddy, Matt Cotten, Jonathan Dambrozio, Simon Dellicour, Antonino Di Caro, Joseph W. Diclaro, Sophie Duraffour, Mike Elmore, Lawrence Fakoli, Merle Gilbert, Sahr M. Gevao, Stephen Gire, Adrianne Gladden-Young, Andreas Gnirke, Augustine Goba, Donald S. Grant, Bart Haagmans, Julian A. Hiscox, Umaru Jah, Brima Kargbo, Jeffrey Kugelman, Di Liu, Jia Lu, Christine M. Malboeuf, Suzanne Mate, David A. Matthews, Christian B. Matranga, Luke Meredith, James Qu, Joshua Quick, Susan D. Pas, My VT Phan, Georgios Poliakis, Chantal Reusken, Mariano Sanchez-Lockhart, Stephen F. Schaffner, John S. Schieffelin, Rachel S. Sealfon, Etienne Simon-Loriere, Saskia L. Smits, Kilian Stoecker, Lucy Thorne, Ekaete A. Tobin, Mohamed A. Vandi, Simon J. Watson, Kendra West, Shannon Whitmer, Michael R. Wiley, Sarah M. Winnicki, Shirlee Wohl, Roman Wölfel, Nathan L. Yozwiak, Kristian G. Andersen, Sylvia Blyden, Fatorma Bolay, Miles Carroll, Bernice Dahn, Boubacar Diallo, Pierre Formenty, Christophe Fraser, George F. Gao, Robert F. Garry, Ian Goodfellow, Stephan Günther, Christian Happi, Edward C Holmes, Brima Kargbo, Paul Kellam, Marion P.G. Koopmans, Nicholas J. Loman, N’Faly Magassouba, Dhamari Naidoo, Stuart T. Nichol, Tolbert Nyenswah, Gustavo Palacios, Oliver G Pybus, Pardis Sabeti, Amadou Sall, Keïta Sakoba, Ute Ströeher, Isatta Wurie, Marc A Suchard, Philippe Lemey, Andrew Rambaut

**Author notes:** The findings and conclusions in this report are those of the authors and do not necessarily represent the official position of the Centers for Disease Control and Prevention (Shannon Whitmer, Stuart T. Nichol, Ute Ströeher).

## Abstract

The 2013-2016 epidemic of Ebola virus disease in West Africa was of unprecedented magnitude, duration and impact. Extensive collaborative sequencing projects have produced a large collection of over 1600 Ebola virus genomes, representing over 5% of known cases, unmatched for any single human epidemic. In a comprehensive analysis of this entire dataset, we reconstruct in detail the history of migration, proliferation and decline of Ebola virus throughout the region. We test the association of geography, climate, administrative boundaries, demography and culture with viral movement among 56 administrative regions. Our results show that during the outbreak viral lineages moved according to a classic ‘gravity’ model, with more intense migration between larger and more proximate population centers. Notably, we find that despite a strong attenuation of international dispersal after border closures, localized cross-border transmission beforehand had already set the seeds for an international epidemic, rendering these measures relatively ineffective in curbing the epidemic. We use this empirical evidence to address why the epidemic did not spread into neighboring countries, showing that although these regions were susceptible to developing significant outbreaks, they were also at lower risk of viral introductions. Finally, viral genome sequence data uniquely reveals this large epidemic to be a heterogeneous and spatially dissociated collection of transmission clusters of varying size, duration and connectivity. These insights will help inform approaches to intervention in such epidemics in the future.

## Main text

At least 28,000 cases and 11,000 deaths (World Health Organization, 2016a) have been attributed to the Makona variant of Ebola virus (EBOV) (Kuhn et al., 2014) in the two and a half years that it circulated in West Africa. The epidemic is thought to have begun in December 2013 in Guinea, but was not detected and reported until March 2014 (Baize et al., 2014). Initial efforts to control the outbreak in Guinea were considered to be succeeding (World Health Organization Regional Office for Africa, 2014), but in early 2014 the virus crossed international borders into neighbouring Liberia (first cases diagnosed in late March) and Sierra Leone (first documented case in late February (Goba et al., 2016; Sack et al., 2014), first diagnosed cases in May (Gire et al., 2014)). Viral genomes sequenced from three patients in Guinea early in the epidemic (Baize et al., 2014) helped to establish that the progenitor of the Makona variant originated in Central Africa and arrived in West Africa within the last 15 years (Dudas and Rambaut, 2014; Gire et al., 2014). Rapid sequencing of the first reported cases in Sierra Leone confirmed that EBOV had crossed the border from Guinea and they were not the result of an independent zoonotic introduction (Gire et al., 2014). Subsequent studies analysed the genetic makeup of the Makona variant but focused on infections in either Guinea (Carroll et al., 2015; Quick et al., 2016; Simon-Loriere et al., 2015), Sierra Leone (Arias et al., 2016; Park et al., 2015) or Liberia (Kugelman et al., 2015; Ladner et al., 2015), identifying local viral lineages and patterns of transmission within each country.

Although virus sequencing has covered considerable fractions of the epidemic in each country, individual studies focused on either limited geographical areas or periods of time, so that the regional level patterns and drivers of the epidemic across its entire duration have remained uncertain. Using 1610 genome sequences collected throughout the epidemic, which represent over 5% of known Ebola virus disease (EVD) cases (Figures 1 & S1), we apply phylogenetic approaches to reconstruct a detailed history of the movement of the virus within and among the three most affected countries. Using a recently developed approach for integrating covariates of spatial spread within a phylogeographic model (Lemey et al., 2014), we test which features of each region (administrative, economic, climatic, infrastructural and demographic) were important in shaping the spatial dynamics of EBOV. We also examine the effectiveness of international border closures on controlling virus dissemination. Finally, we investigate why regions that immediately border the most affected countries did not develop protracted outbreaks similar to those that ravaged Sierra Leone, Guinea and Liberia.

**Figure 1.**
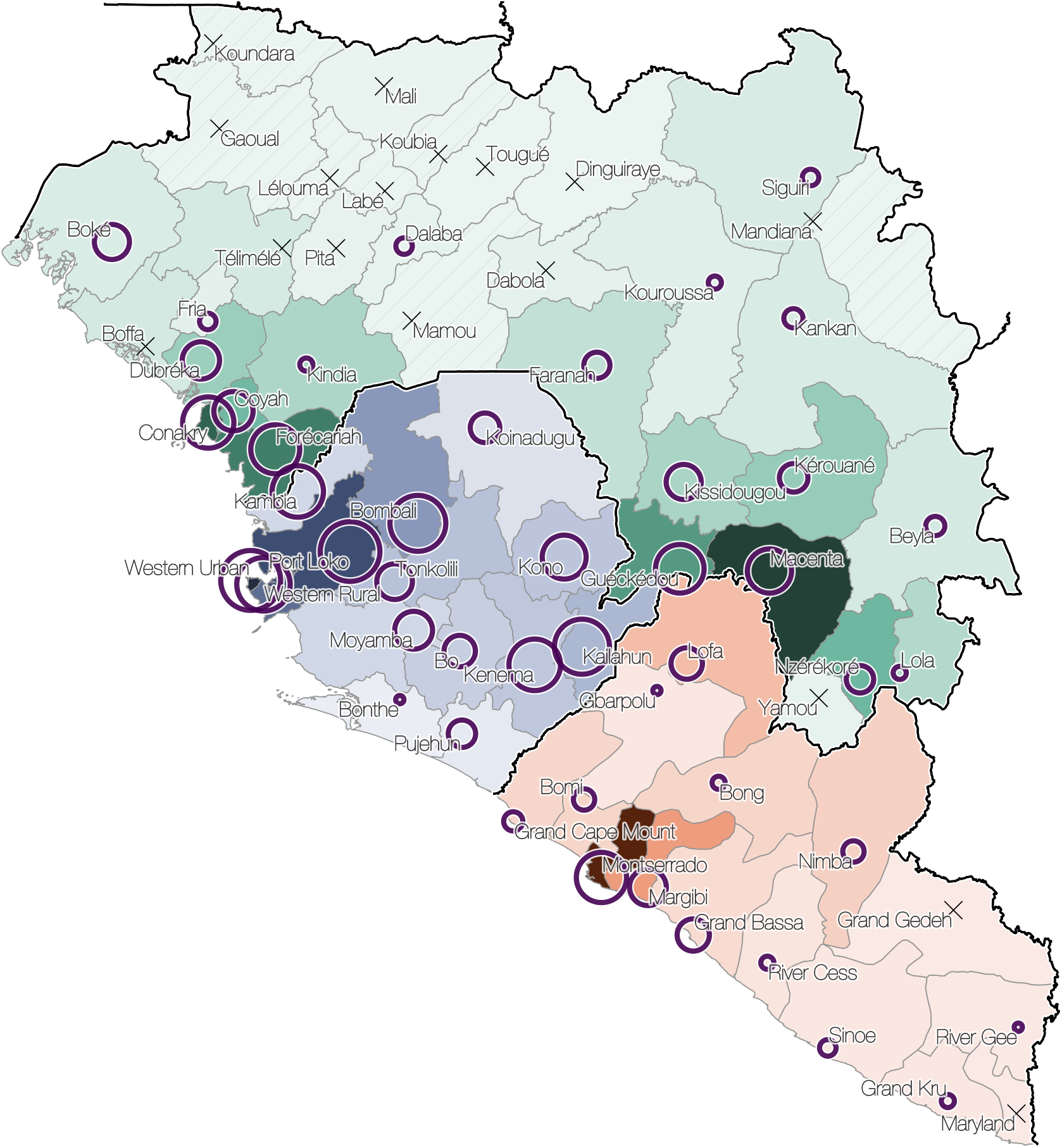
Distribution of EBV cases and virus sequences. Administrative regions within Guinea (green), Sierra Leone (blue) and Liberia (red); shading is proportional to the cumulative number of known and suspected EVD cases in each region. Darkest shades represent 784 cases for Guinea (Macenta), 3219 cases for Sierra Leone (Western Urban) and 2925 cases for Liberia (Montserrado); hatched areas indicate regions without any reported EVD cases. Circle diameters are proportional to the number of sequences available from that region over the entire epidemic with the largest circle representing 152 sequences. Crosses mark regions for which no sequences are available. Circles and crosses are positioned at population centroids within each region. The number of sequences and number of cases for each region where cases were recorded are strongly correlated (Spearman rank correlation coe cient 0.93; Supplementary Figure S1).

## Origin, ignition and trajectory of the epidemic

Molecular clock dating indicates that the most recent common ancestor of the epidemic existed in early December 2013 (95% highest posterior density interval: Oct 2013, Feb 2014) and phylogeographic estimation assigns this ancestor to the Guéckéedou préefecture with a high degree of confidence (96% posterior support) (Figure 2). In addition, we find that initial lineages deriving from this common ancestor circulated among Guéckéedou and its neighbouring préefectures of Macenta and Kissidougou until late February 2014 (Figure 2). These results, based on a comprehensive sample of EBOV genomes, support the epidemiological evidence that the West African epidemic began in late 2013 in Guéckédou préfecture of Guinea (Baize et al., 2014).

**Figure 2.**
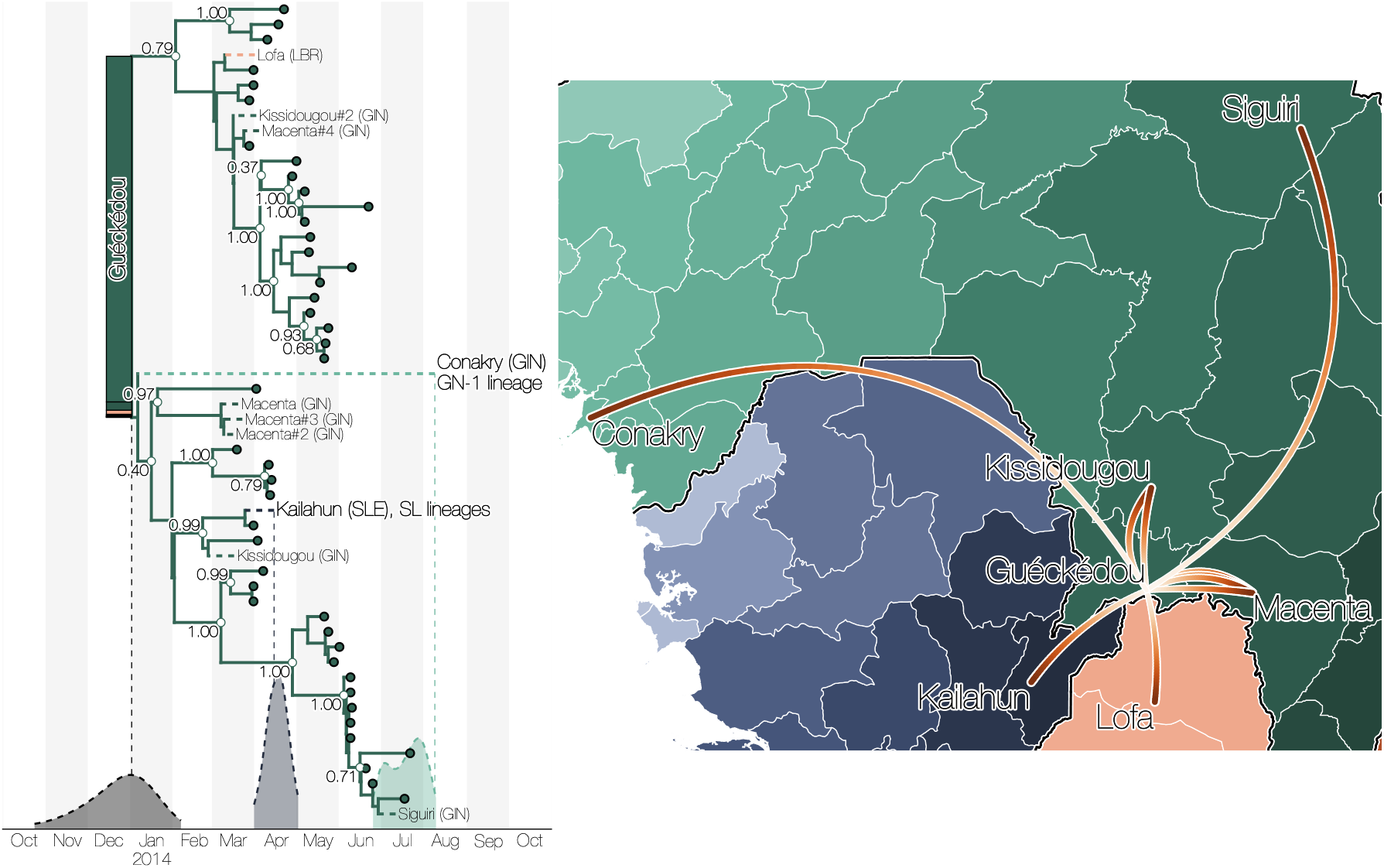
Summary of early epidemic events. a) The time-scaled phylogeny of the early sampled cases in Guéckédou, Guinea and their relationships to the initial dispersal events into other neighbouring and more distant regions. Stacked bars at the root of the tree indicate posterior probabilities for the origin of the epidemic (0.96 for Guéckédou, 0.02 for Macenta, 0.01 for Lofa and negligible probabilities for other locations). 95% posterior densities of the time of the common ancestor of all lineages (grey) and far-dispersing lineages into Kailahun district (blue, introduction gave rise to SL lineages) and to Conakry préfecture (green, introduction leads to lineage GN-1) are shown at the bottom of the tree. Nodes with three or more tips have posterior probabilities shown if >0.3. b) These same dispersal events (marked by dashed lineages on the phylogeny) projected on a map with directionality indicated by colour intensity (from white to red). Lineages that migrated to Conakry and Kailahun have led to the vast majority of cases throughout the region.

The first introduction of EBOV from Guinea into another country that resulted in sustained transmission is estimated to have occurred in early April 2014 (Figure 2), when the virus spread to the Kailahun district of Sierra Leone (Goba et al., 2016; Sack et al., 2014). This lineage was first detected in Kailahun at the end of May 2014, from where it spread across the region (Figure 3 & S2). From Kailahun EBOV spread extremely rapidly in May 2014 into several counties of Liberia (Lofa, Montserrado and Margibi) (Ladner et al., 2015) and Guinea (Conakry, back into Guéckédou) (Carroll et al., 2015; Simon-Loriere et al., 2015). The virus continued to spread westwards through Sierra Leone, and by July 2014 it was present in the capital city, Freetown.

**Figure 3.**
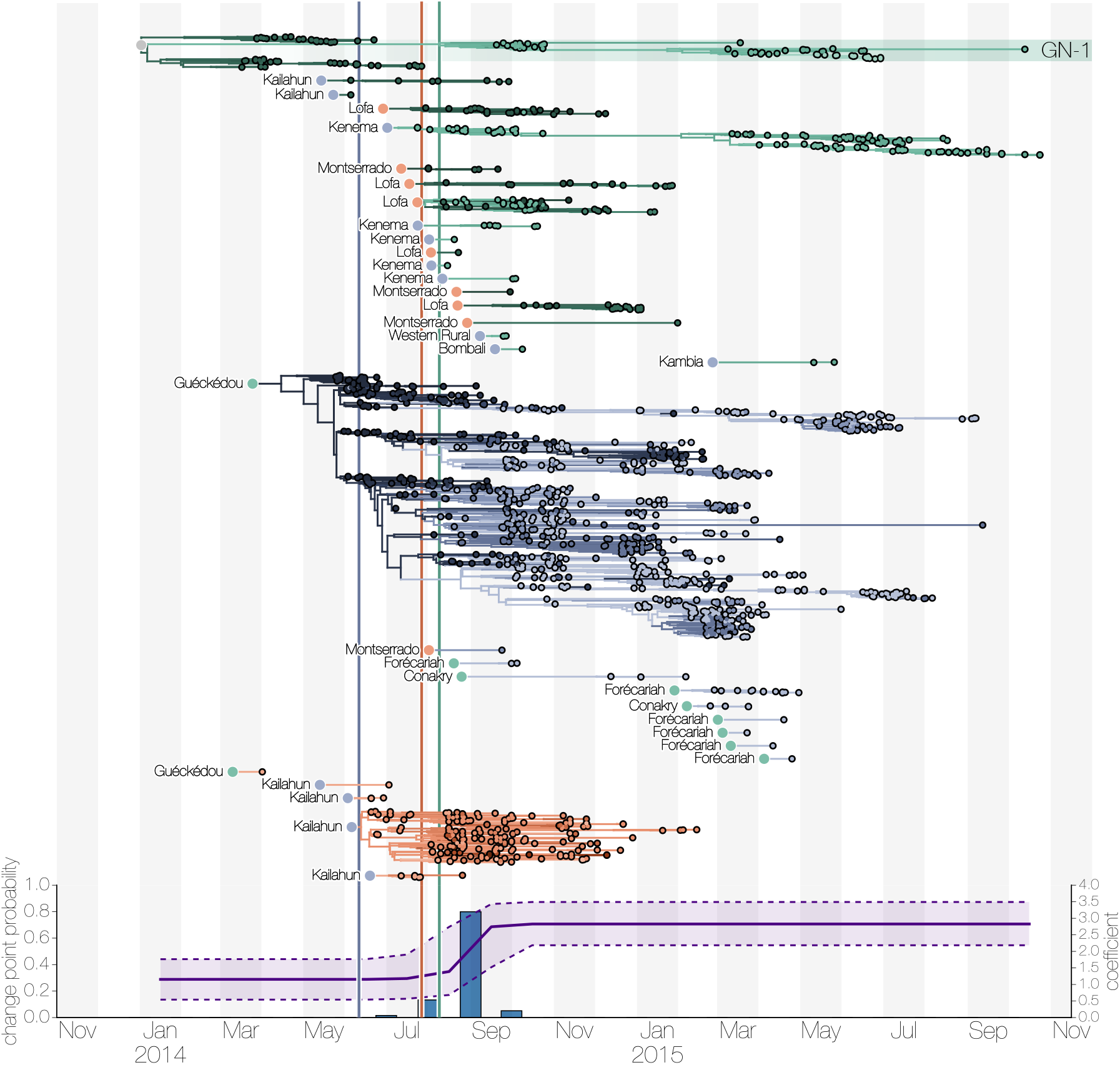
Time-scaled phylogeny deconstructed into country-specific transmission chains arising from independent international movements. a) EBOV lineages, tracked until the sampling date of their last known descendants, sorted by country (Guinea, green; Sierra Leone, blue; Liberia, red) and earliest possible introduction date. Tips are shared by longitude (lightest to the West, darkest to the East). Circles at root of each subtree denote the country of origin for the introduced lineage. The four introductions into Liberia in May-June 2014 are all inferred to have come from Kailahun, Sierra Leone. These may represent a few, or just one, movement events; however, the genetic similarity of these Liberian genomes to viruses from Kailahun makes further resolution impossible. In contrast, the multiple introductions into Guinea are very likely the result of multiple separate movement events over a 10 month period. b) Epoch estimates of the change point probability (primary Y-axis) and log coe cient (mean and credible interval; secondary Y-axis) for the within country-effect (the only effect with support for epoch dynamics; see Supplementary Information). The highest change point probability and an associated doubling of log effect size for within country transmission is estimated between August and September 2014 (blue columns). Vertical lines represent dates of border closures by the respective countries (Sierra Leone, blue; Liberia, red; Guinea, green).

By mid-September 2014 Liberia was reporting >500 new EVD cases per week, mostly driven by a large outbreak in Montserrado county, which encompasses the capital city, Monrovia. Sierra Leone reported as many as 700 new cases per week by mid-November, driven by large outbreaks in Port Loko, Western Urban (Freetown) and Western Rural districts (Freetown suburbs). December 2014 brought the first signs that efforts to control the epidemic in Sierra Leone were effective as EVD incidence began dropping. By March 2015 the epidemic was largely under control in Liberia and eastern Guinea, although sustained transmission was still occurring in western Guinea and western Sierra Leone, near the border between the two countries. By the following month prevalence had declined such that only a handful of relatively distantly related lineages survived from the exponential growth phase of the epidemic (Arias et al., 2016; Quick et al., 2016) (Figure 3).

The last Ebola virus resulting from a conventionally-acquired infection was collected and sequenced in October 2015 in Forecariah préfecture (Guinea) (Quick et al., 2016). Following this, only sporadic cases of EVD were detected: in Margibi (Liberia) in June 2015, Montserrado (Liberia) in November 2015, Tonkolili (Sierra Leone) in January and February 2016, and Nzérékoré (Guinea) in March 2016. All these sporadic cases likely result from transmission from EVD survivors with established persistent infections (Blackley et al., 2016; Mate et al., 2015).

## Factors associated with EBOV dispersal

To determine the factors that influenced the spread of EBOV among administrative regions at the district (Sierra Leone), préfecture (Guinea) and county (Liberia) levels we employed a phylogeographic generalized linear model (GLM) (Lemey et al., 2014). Of the 25 factors assessed (see Table S2 for a full list and description) five were included in the model with categorical support (Table 1). In summary, EBOV migration events tend to occur between geographically close regions (great circle distance: Bayes factor (BF) support for inclusion BF>50). Half of all virus lineage movements occurred between locations <72 km apart and only 5% involved movement over 232 km (Figure 5a). Population sizes are very strongly (BF>50) positively correlated with viral dissemination, with a stronger effect for the population size of the origin location than that for the destination population size. The result, when combined with the inverse effect of geographic distance, implies that the epidemic's spread followed a classic gravity-model dynamic. Gravity models, widely used in economic and geographic studies, describe the movement of people between locations as a function of their population sizes and distance apart. They are a natural choice for modelling infectious disease transmission (Truscott and Ferguson, 2012; Viboud et al., 2006) and have been used in spatio-temporal modelling of EBOV transmission in Sierra Leone (Yang et al., 2015). Here we use viral genomes to provide empirical evidence that such a process drove viral dissemination during the epidemic.

**Table 1.**
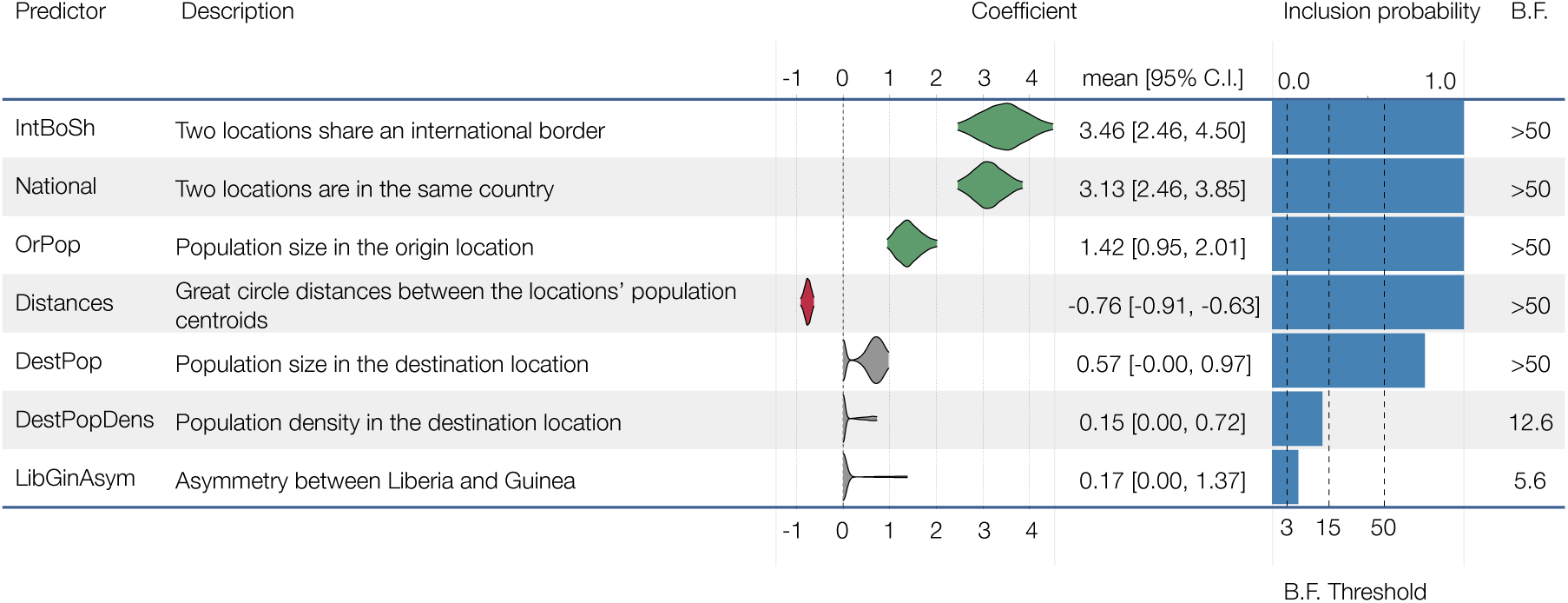
Summary of the phylogenetic generalized linear model results. The estimated coe cients and model inclusion probabilities for spatial movement predictors supported with a Bayes factor (BF) >3. Positive coe cients are shown in green, negative in red. The remainder are not supported and are not shown (see supplementary document for a full list).

In addition to geographical distance, we found a significant propensity for migration events to occur among administrative regions within each country, as opposed to international viral dispersal (National effect, BF>50), suggesting that country borders acted to curb the geographic spread of EBOV. Within-country viral migration is higher than international movement even after the direct effect of distance is accounted for. When international migrations do take place, they are more intense between administrative regions that meet on an international border (IntBoSh, BF>50).

We also tested whether sharing of any of 17 vernacular languages explains virus spread, as this might that reflect local cultural links including those between non-contiguous or international regions, but we found no evidence that such linguistic links were correlated with EBOV spread. A variety of other variables that might intuitively contribute to EBOV transmission, such as aspects of urbanization (economic output, population density, travel times to large settlements) and climatic effects were not found to be significantly associated with EBOV migration. However, these factors may have contributed to the size and longevity of outbreaks after their introduction to a region (see below).

## Factors associated with local EBOV proliferation

The analysis above identified factors that predict virus movement between administrative regions. These factors represent the degree of importation risk, i.e. the likelihood that a viral lineage initiates at least one infection in a new region, and are dominated by geographical and administrative factors. However, the epidemiological consequences of each introduction — the size and duration of resulting transmission chains — may be affected by different factors. To investigate this we explored which demographic, economic and climatic factors might predict cumulative case counts (World Health Organization, 2016a) for each region (Bayesian generalized linear model; see Supplementary Methods).

We find that cumulative case counts in each location were associated with factors related to urbanization (Table 2): primarily population sizes (PopSize, BF 29.6) and a significant inverse association with travel times to the nearest settlement with >50,000 inhabitants (tt50K, BF 32.4). These results confirm the common perception that, compared to previous EVD outbreaks, widespread transmission within urban regions in West Africa was a major contributing factor to the scale of the epidemic of the Makona variant.

**Table 2.**
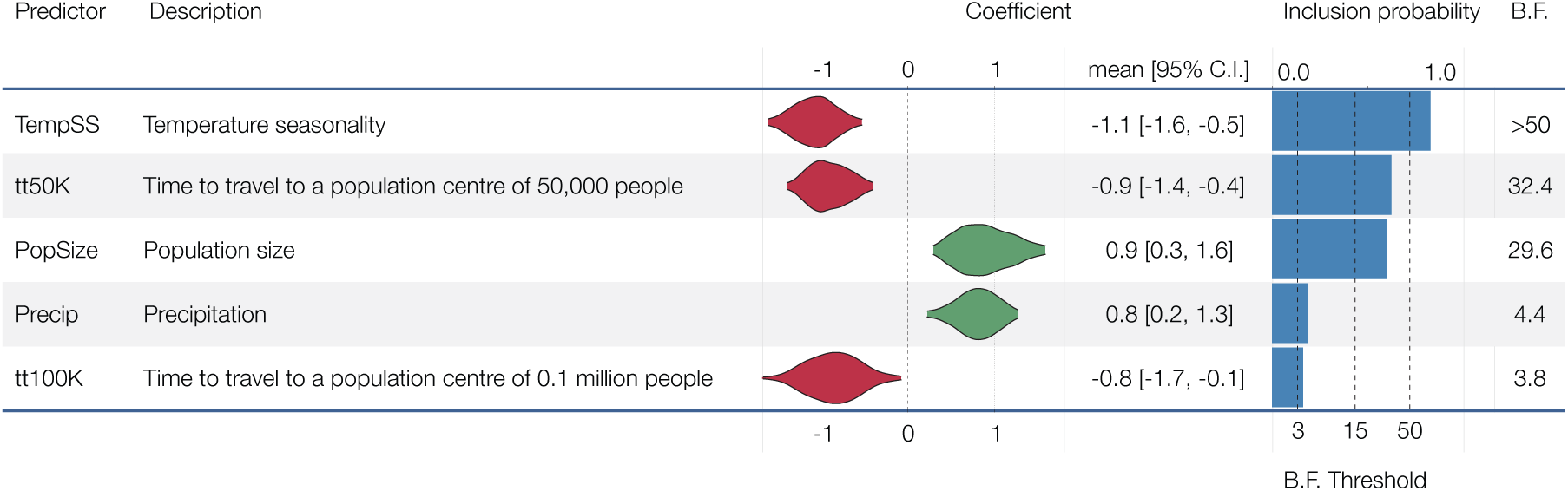
Summary of generalized linear model results with case counts as the response variable. The estimated coe cients and model inclusion probabilities for per-region predictors supported with a Bayes factor (BF) >3. Positive coe cients are shown in green, negative in red. The remainder are not supported and are not shown (see supplementary document for a full list).

As the epidemic in West Africa progressed there were fears that increased rainfall and humidity might make the Ebola virus more environmentally stable, especially in light of frequent post-mortem transmission of the virus (Fischer et al., 2015). Although we found no evidence of an association between EBOV migration and any aspects of local climate, we find that regions with less seasonal variation in temperature, and more rainfall, tended to have larger EVD outbreaks (TempSS, BF >50 and Precip, BF 4.4 respectively).

## Did international travel restrictions have an effect?

It has been suggested that porous borders between Liberia, Sierra Leone and Guinea allowed unimpeded spread of EBOV during the 2013-2016 epidemic (Bausch and Schwarz, 2014; Chan, 2014; Wesolowski et al., 2014). Our results suggest that, on average, international borders were associated with a decreased rate of transmission events compared to national borders (Figure S3), but there were still frequent international cross-border transmission events. Specifically, these events were concentrated in Guéckédou (Guinea), Kailahun (Sierra Leone) and Lofa (Liberia) during the early phases of the epidemic (Figure S4b), and in the later stage of the epidemic (Figure S4b) between neighbouring Forécariah (Guinea) and Kambia (Sierra Leone). These later movements significantly hindered efforts to interrupt the final chains of transmission in late 2015, with a number of such chains moving back and forth across this border (Arias et al., 2016; Goodfellow et al., 2015; Quick et al., 2016). Sierra Leone announced border closures on 11 June 2014, followed by Liberia on 27 July 2014, and Guinea on 9 August 2014, although there is little information on what these border closures actually entailed. As a consequence, even though our results show that international viral spread was more intense before these borders were closed (mean change point: Aug-Sept 2014; 80.0% posterior support; (Figure 3b; see also Figure S5), it is difficult to ascertain whether it was the border closures themselves that were responsible for the apparent reduction in cross-border transmissions, as opposed to concomitant control efforts or public information campaigns. Overall, these results suggest that border closures may have reduced international traffic, particularly over longer distances and between larger population centres, but by the time Sierra Leone and later Liberia closed their borders the epidemic had become firmly established in both countries (Figure 3).

## Why did the epidemic not spread further?

With the exception of a few documented exportations (Abdoulaye et al., 2015; Folarin et al., 2016; Hoenen et al., 2015), the West African Ebola virus epidemic did not spread into neighbouring regions of Guinea-Bissau, Senegal, Mali, or Côte d’Ivoire, and no cases were reported in seven préfectures of northern Guinea. By extending our GLM (i.e., the supported predictors and their estimated coefficients) to include these regions we can address whether these regions were spared EBOV cases through good fortune, or because they had an inherently lower risk of EBOV spread and transmission. We estimated the degree to which these, apparently EVD-free, regions had the potential to be exposed to viral introductions from regions with cases (see supplementary methods). Overall, the contiguous regions in neighbouring countries were all predicted to low numbers of introductions (Figure 4a). They were not, however, predicted to have particularly low levels of transmission if an outbreak had been seeded (Figure 4b). Thus, it is likely that some of these surrounding regions and their countries overall were at risk of an EVD epidemic, but that their geographical distance from areas of active transmission and the attenuating effect of international borders prevented their epidemic potential from being realized. The Kati region in Mali and Tonkpi region in Côte d’Ivoire are to some extent exceptions to this general result, being more susceptible to viral introductions under the gravity model because of their large populations (Kati, 1 million; Tonkpi 950,000), (Figure 4a) and are predicted to have experienced many cases had EVD become established (Figure 4b).

**Figure 4.**
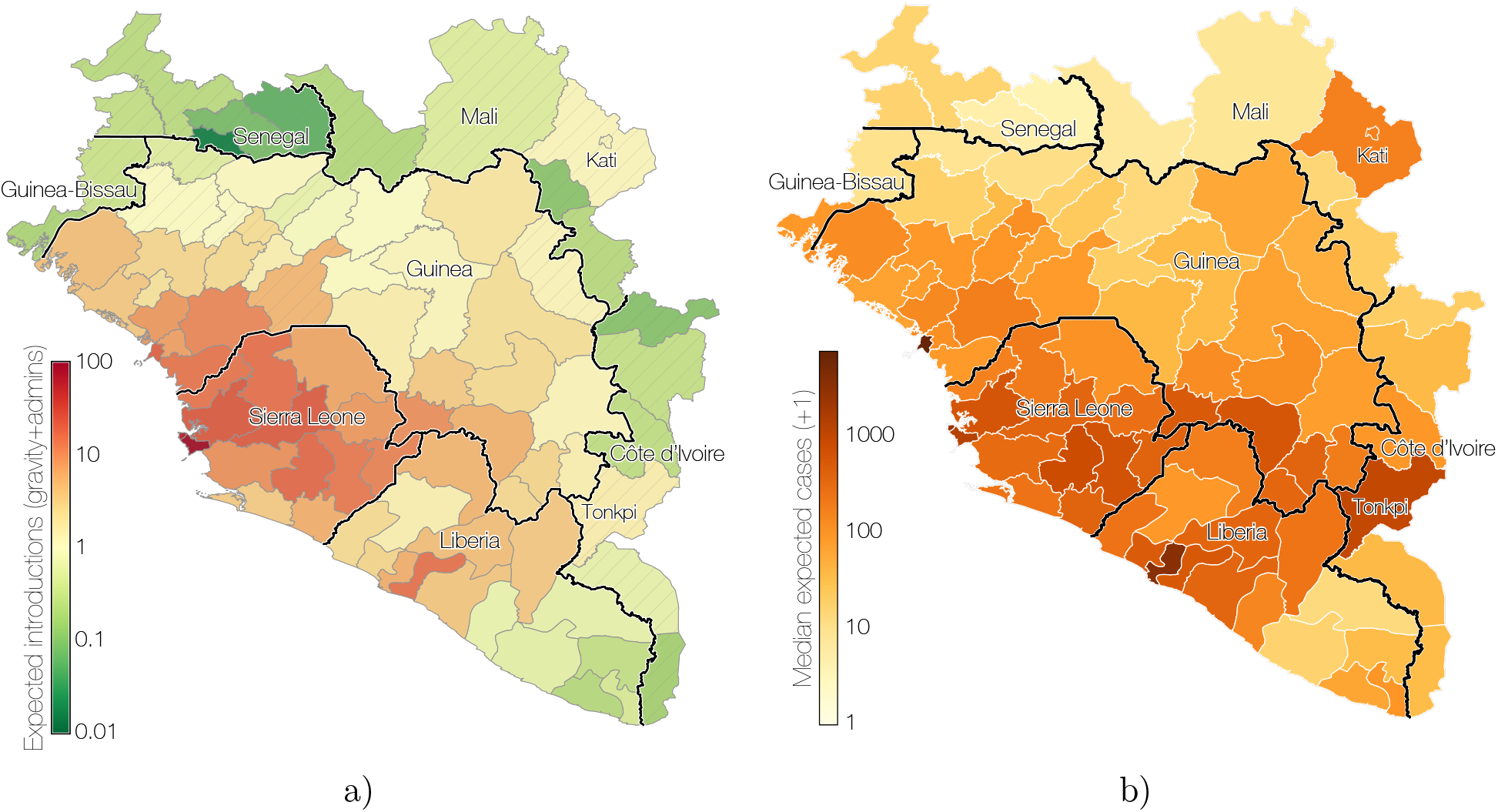
Predicted destinations and consequences of viral migrations. a) Predicted number of imports into each of 63 regions in Guinea, Sierra Leone and Liberia (including 7 with no recorded cases in Guinea) and the surrounding 18 regions from the neighbouring countries of Guinea-Bissau, Senegal, Mali and Côte d’Ivoire. The expected number of exports from locations in the phylogeographic tree and imports to any location are calculated based on the phylogeographic GLM model estimates and associated predictors that were extended to apparently EVD-free locations (see Supplementary Methods). b) Predicted outbreak sizes from the generalized linear model fitted to case data.

## The metapopulation structure and dynamics of the epidemic

Figure 3 shows that after the initial establishment of transmission in Sierra Leone and Liberia, Guinea experienced repeated reintroductions of viral lineages from the escalating epidemics in these other two countries. From the 5% of cases that were sequenced, our analysis reveals that there were at least 21 (95% credible interval, CI: 18 - 24) re-introductions into Guinea from April 2014 to February 2015. Although an early epidemic lineage was established in the region around the Guinean capital, Conakry, and persisted for the duration of the epidemic (GN-1 in Figures 2 & 3), the continual ‘seeding’ of EBOV lineages into Guinea without a clear peak in transmission suggests that the virus may have been struggling to maintain transmission in that country. There were also numerous introductions into Sierra Leone over a similar time period (median: 9, 95% CI: 7 - 11) but the resulting transmission chains constituted a tiny proportion of the Sierra Leonean epidemic, with the bulk of transmission resulting from one early introduction (Figure 3a).

The importance of repeated seeding as a factor in the longevity of the epidemic is also suggested by the pattern of viral movement among administrative regions within each country (Figure S6). Regional epidemics were the result of multiple overlapping introduction events followed by within-region spread and occasional onward transmission to other regions. This observation suggests a metapopulation model in which viral persistence is driven by introduction into novel contact networks rather than by mass-action susceptible-infectious-recovered (SIR) dynamics (Ferrari et al., 2008). We find that, on average, EBOV migrates between administrative regions at a rate of 0.85 events per lineage per year (95% CI: 0.72, 0.97). If we assume a serial interval of 15.3 days (WHO Ebola Response Team, 2014), this translates to a 3.6% chance (95% CI: 3.0%, 4.1%) that a single step in the transmission chain migrates between regions. The detection and isolation of these mobile cases may have a disproportionate effect on the control of the epidemic.

Many regions experienced numerous independent introductions (Figure 5b) but the size of the clusters of cases that result from these introductions was generally small (with a mean cluster size of 4.3 and only 5% larger than 17 in our sample; Figure 5c) and their persistence of limited duration (a mean persistence time of 41.3 days with only 5% greater than 181 days; Figure 5d). Here, we define a ‘cluster’ as a group of sequenced cases that derive from a single introduction event into a region without including subsequent infections in other regions and persistence as the time between the introduction event and the last sampled case in the cluster. These definitions are conservative with regards to sampling intensity as we expect additional samples would split apart clusters rather than join them. Furthermore, introductions that were not detected will be disproportionately smaller, and so the cluster size estimate will be biased towards larger sizes. Thus, with 5.8% sampling, we arrive at a conservative estimate of approximately 75 regional cases per introduction event. Although larger population centres, in particular the capital cities, generally had more introductions (Figure S7a) the cluster sizes are less strongly associated with population size (Figure S7b). The frequent extinction of these clusters even though a small fraction of individuals were infected suggests that they were constrained by the degree of connectedness among contact networks. Thus, it appears the West African epidemic was sustained by frequent seeding that resulted in numerous small local clusters of cases, some of which went on to seed further local clusters.

**Figure 5.**
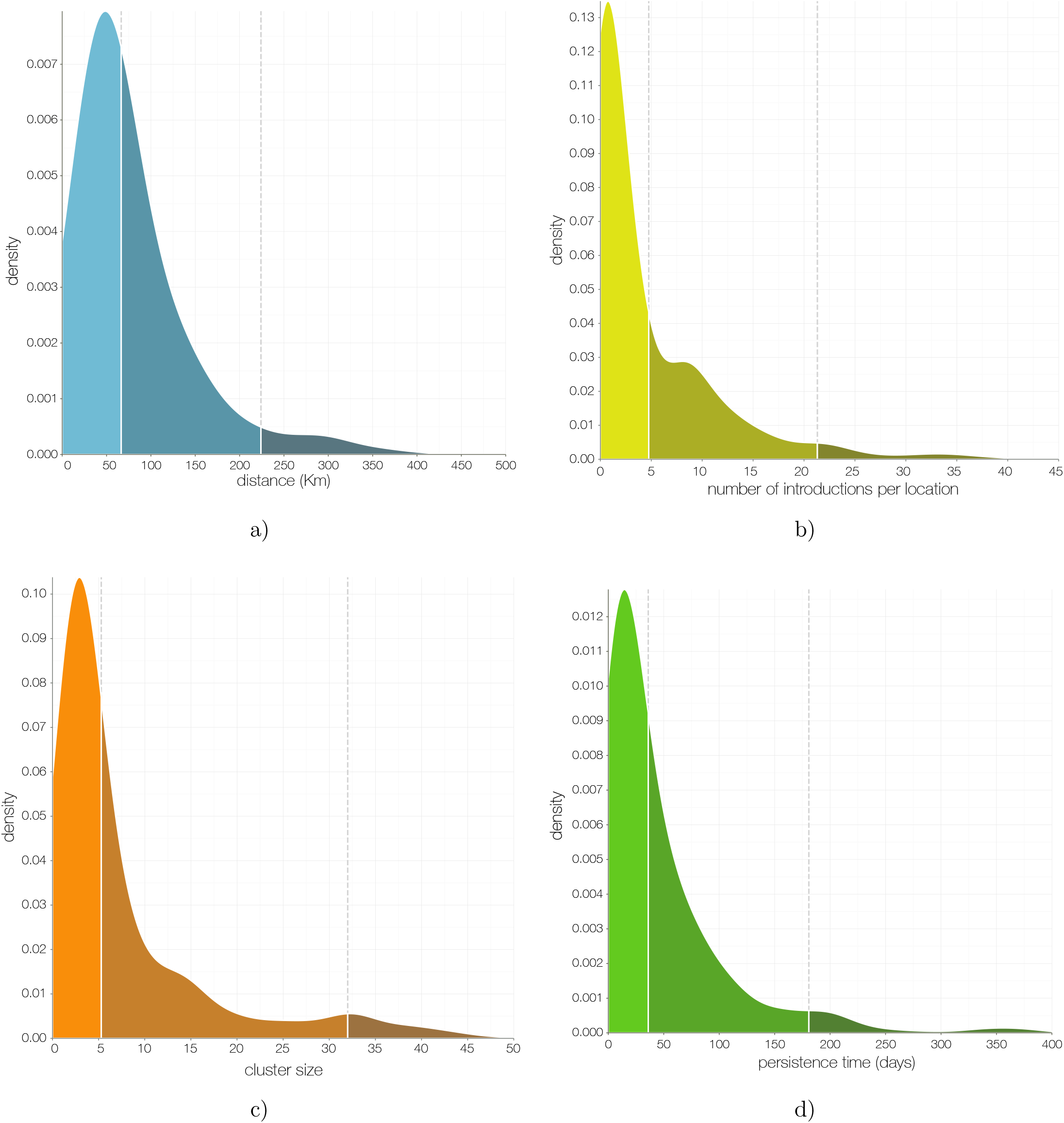
The metapopulation structure of the epidemic. a) Kernel density estimate (KDE) of distance for all inferred migrations: 50% occur over distances <72 km and <5% occur over distances >232 km. b) KDE of the number of independent introductions into each administrative region: 50% have fewer than 4.8 and <5% greater than 21.3. c) KDE of the mean size of sampled cases resulting from each introduction with at least 2 sampled cases: 50% <5.3, 95% <32. d) KDE of the persistence of clusters in days (from time of introduction to time of the last sampled case): 50% <36 days, 95% <181 days.

## Viral genomics as a tool for outbreak response

The 2013-2016 EVD epidemic in West Africa has unfortunately become a costly lesson in dealing with an infectious disease outbreak when both the exposed population and the international community are unprepared. It also demonstrates the value of pathogen genome sequencing in a public healthcare emergency situation and the value of timely pre-publication data sharing in order to identify the origins of imported lineages, to track viral transmission as the epidemic progresses, and to follow up on individual cases as the epidemic subsides. Real-time virus genome sequencing at the point of diagnosis can provide additional insights, especially when conventional epidemiological contact tracing is challenging. Other sources of human mobility data, mobile phone network data in particular, are promising but currently such data is difficult to obtain in a timely fashion (Wesolowski et al., 2014). It is inevitable that as sequencing becomes cheaper, more portable and accurate, real-time viral surveillance and molecular epidemiology will be routinely deployed on the frontlines of infectious disease outbreaks (Gardy et al., 2015; Yozwiak et al., 2015; Woolhouse et al., 2015; Quick et al., 2016). As viral genome sequencing is scaled up and gets closer to the time-scale of viral evolution, the pressure will increasingly fall on analysis techniques to provide the necessary temporal resolution to inform outbreak response. The analysis of the comprehensive EBOV genome set collected during the 2013-2016 epidemic, including the findings presented here and in other studies (Arias et al., 2016; Carroll et al., 2015; Gire et al., 2014; Kugelman et al., 2015; Ladner et al., 2015; Park et al., 2015; Simon-Loriere et al., 2015; Stadler et al., 2014; Tong et al., 2015) will provide a framework for predicting the behaviour of future outbreaks for EBOV, other filoviruses, and perhaps other human pathogens.

Many open questions remain about the biology of EBOV. As sustained human-to-human transmission waned, West Africa experienced several instances of recrudescent transmission, often in regions that had not seen cases for many months as a result of persistent sub-clinical infections (Blackley et al., 2016; Mate et al., 2015; World Health Organization, 2016c,b). Although, in hindsight, such sequelae were not entirely unexpected (Rowe et al., 1999), the magnitude of the 2013-2016 epidemic has put the region at ongoing risk of sporadic EVD re-emergence. Similarly, the nature of the reservoir of EBOV, and its geographic distribution, remain as fundamental gaps in our knowledge. Resolving these questions is critical to predicting the risk of zoonotic transmission and hence of future outbreaks of this devastating disease.

## Methods Summary

A total of 1610 nearly complete EBOV genome sequences were collated, aligned and annotated with date of sampling and likely location of infection (all data available from https://github.com/ebov/space-time). Geographical, demographic and climatic variables were collated for each of 63 regions in three focal countries, and for a further 18 regions in surrounding countries that reported no cases or no sustained transmission (see supplemental information for details). Time structured phylogenies were inferred using BEAST (Drummond et al., 2012; Ayres et al., 2012) and these formed the basis of a phylogenetic generalized linear model (Lemey et al., 2014) that infers the probability of inclusion, and degree of correlation, of each of the predictor variables for the spatial pattern of virus lineage migration. Along each branch of the tree we infer change among regions (Minin and Suchard, 2008). For those variables in the model with significant support, we extended the analysis to allow a single step-change in coefficient and inferred the time of this change-point. Furthermore, we used the inferred spatial model to estimate the expected number of migrations into regions which experience no known cases of EVD including in the surrounding countries. Finally, to assess which of the demographic and climatic variable were predictive of the magnitude of outbreak once introduced into a region, we employed generalized linear models and Bayesian model averaging, with cumulative case counts in each affected region as a response variable.

## Supplementary Methods

### Sequence data

We compiled a data set of 1610 publicly available full Ebola virus (EBOV) genomes sampled between 17 March 2014 and 24 October 2015 (see https://github/ebov/space-time/data/ for full list and metadata). The number of sequences and the proportion of cases sequenced varies with country; our data set contains 209 sequences from Liberia (3.8% of known and suspected cases), 982 from Sierra Leone (8.0%) and 368 from Guinea (9.2%) (Table S1). Most (1100) genomes are of high quality, with ambiguous sites and gaps comprising less than 1% of total alignment length, followed by sequences with between 1% and 2% of sites comprised of ambiguous bases or gaps (266), 98 sequences with 2-5%, 120 sequences with 5-10% and 26 sequences with more than 10% of sites that are ambiguous or are gaps. Sequences known to be associated with sexual transmission or latent infections were excluded, as these viruses often exhibit anomalous molecular clock signals (Blackley et al., 2016; Mate et al., 2015). Sequences were aligned using MAFFT (Katoh et al., 2002) and edited manually. The alignment was partitioned into coding regions and non-coding intergenic regions with a final alignment length of 18992 nucleotides (available from https://github/ebov/space-time/data/).

### Masking putative ADAR edited sites

As noticed by Tong et al. (2015), Park et al. (2015) and other studies, some EBOV isolates contain clusters of T-to-C mutations within relatively short stretches of the genome. Interferon-inducible adenosine deaminases acting on RNA (ADAR) are known to induce adenosine to inosine hypermutations in double-stranded RNA (Bass and Weintraub, 1988). ADARs have been suggested to act on RNAs from numerous groups of viruses (Gélinas et al., 2011). When negative sense single stranded RNA virus genomes are edited by ADARs, A-to-G hypermutations seem to preferentially occur on the negative strand, which results in U/T-to-C mutations on the positive strand (Cattaneo et al., 1988; Rueda et al., 1994; Carpenter et al., 2009). Multiple T-to-C mutations are introduced simultaneously via ADAR-mediated RNA editing which would interfere with molecular clock estimates and, by extension, the tree topology. We thus designate four or more T-to-C mutations within 300 nucleotides of each other as a putative hypermutation tract, whenever there is evidence that all T-to-C mutations within such stretches were introduced at the same time, *i.e.* every T-to-C mutation in a stretch occurred on a single branch. We detect a total of 15 hypermutation patterns with up to 13 T-to-C mutations within 35 to 145 nucleotides. Of these patterns, 11 are unique to a single genome and 4 are shared across multiple isolates, suggesting that occasionally viruses survive hypermutation are transmitted (Smits et al., 2015). Putative tracts of T-to-C hypermutation almost exclusively occur within non-coding intergenic regions, where their effects on viral fitness are presumably minimal. In each case we mask out these sites as ambiguous nucleotides but leave the first T-to-C mutation unmasked to provide phylogenetic information on the relatedness of these sequences.

### Phylogenetic inference

Molecular evolution was modelled according to a HKY+Γ_4_ (Hasegawa et al., 1985; Yang, 1994) substitution model independently across four partitions (codon positions 1, 2, 3 and non-coding intergenic regions). Site-specific rates were scaled by relative rates in the four partitions. Evolutionary rates were allowed to vary across the tree according to a relaxed molecular clock that draws branch-specific rates from a log-normal distribution (Drummond et al., 2006). A non-parametric coalescent ‘Skygrid’ tree prior was employed for demographic inference (Gill et al., 2013). The overall evolutionary rate was given an uninformative continuous-time Markov chain (CTMC) reference prior (Ferreira and Suchard, 2008), while the rate multipliers for each partition were given an uninformative uniform prior over their bounds. All other priors used to infer the phylogenetic tree were left at their default values. BEAST XML files are available from https://github/ebov/space-time/data/.

### Geographic history reconstruction

The level of administrative regions within each country was chosen so that population sizes between regions are comparable. For each country the appropriate administrative regions were: préfecture for Guinea (administrative subdivision level 2), county for Liberia (level 1) and district for Sierra Leone (level 2). We refer to them as regions (63 in total but only 56 are recorded to have had EVD cases) and each sequence, where available, was assigned the region where the patient was recorded to have been infected as a discrete trait. When the region within a country was unknown (N=222), we inferred the sequence location as a latent variable with equal prior probability over all available regions within that country. In the absence of any geographic information (N=2) we inferred both the country and the region of a sequence.

We deploy an asymmetric continuous-time Markov chain (CTMC) (Lemey et al., 2009; Edwards et al., 2011) matrix to infer instantaneous transitions between regions. For 56 regions with recorded EVD cases, a total of 3080 independent transition rates would be challenging to infer from one realisation of the process, even when reduced to a sparse migration matrix using stochastic search variable selection (SSVS) (Lemey et al., 2009).

Thus, to infer the spatial phylogenetic diffusion history between the *K* = 56 locations, we adopt a sparse generalized linear model (GLM) formulation of continuous-time Markov chain (CTMC) diffusion (Lemey et al., 2014). This model parameterizes the instantaneous movement rate Λ_*ij*_ from location *i* to location *j* as a log-linear function of *P* potential predictors **X**_*ij*_=(*x_ij_*_1_*, …, x_ijP_*)*t* with unknown coefficients ***β***=(*β*_1_*, …, β_P_*)′ and diagonal matrix ***δ*** with entries (*δ*_1_*, …, δ_P_*). These latter unknown indicators *δ_p_ ∈ {*0, 1} determine predictor *p*'s inclusion in or exclusion from the model. We generalize this formulation here to include two-way random effects that allow for location origin- and destination-specific variability. Our two-way random effects GLM becomes

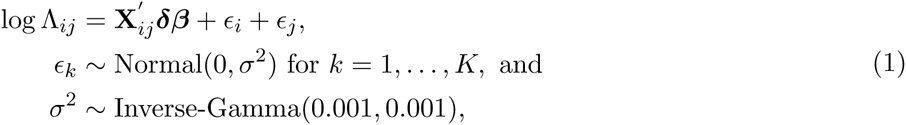

where ***ϵ***=(*ϵ*_1_*, …, ϵ_K_*) are the location-specific effects. These random effects account for unexplained variability in the diffusion process that may otherwise lead to spurious inclusion of predictors.

We follow Lemey et al. (2014) in specifying that *a priori* all *β*_*p*_ are independent and normally distributed with mean 0 and a relatively large variance of 4 and in assigning independent Bernoulli prior probability distributions on *δ_p_*.

Let *q* be the inclusion probability and *w* be the probability of no predictors being included. Then, using the distribution function of a binomial random variable it is straightforward to see that *q*=1 − *w*^1*/P*^, where *P* is the number of predictors, as before. We use a small success probability on each predictor's inclusion that reflects a 50% prior probability (*w*) on no predictors being included.

In our main analysis, we consider 25 individual predictors that can be classified as geographic, administrative, demographic, cultural and climatic covariates of spatial spread (Table S2). Where measures are region-specific (rather than pairwise region measures), we specify both an origin and destination predictor. We also tested for sampling bias by including an additional origin and destination predictor based on the residuals for the regression of sample size against case count (cfr. Fig. S1), but these predictors did not yield any noticeable support (data not shown).

To draw posterior inference, we follow Lemey et al. (2014) integrating ***β*** and ***δ***, and further employ a random-walk Metropolis transition kernel on ***ϵ*** and sample *σ*^2^ directly from its full conditional distribution using Gibbs sampling.

To obtain a joint posterior estimate from this joint genetic and phylogeographic model, two independent MCMC chains were run in BEAST 1.8.4 (Drummond et al., 2012) for 100 million states, sampling every 10 000 states. The first 1000 samples in each chain were removed as burnin, and the remaining 18 000 samples combined between the two runs. These 18 000 samples were used to estimate a maximum clade credibility tree and to estimate posterior densities for individual parameters.

To obtain realisations of the phylogenetic CTMC process, including both transitions (Markov jumps) between states and waiting times (Markov rewards) within states, we employ posterior inference of the complete Markov jump history through time (Minin and Suchard, 2008; Lemey et al., 2014). In addition to transitions ‘within’ the phylogeny, we also estimate the expected number of transitions ‘from’ origin location *i* in the phylogeographic tree to arbitrary ‘destination’ location *j* as follows:

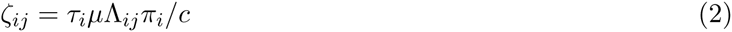

where *τ*_*i*_ is the waiting time (or Markov reward) in ‘origin’ state *i* throughout the phylogeny, *µ* is the overall rate scalar of the location transition process, *π*_*i*_ is the equilibrium frequency of ‘origin’ state *i* and *c* is the normalising constant applied to the CTMC rate matrices in BEAST. To obtain the expected number of transitions to a particular destination location from any phylogeographic location (integrating over all possible locations across the phylogeny), we sum over all 56 origin locations included in the analysis. We note that the destination location can also be a location that was not included in the analysis because we only need to consider destination *j* in the instantaneous movement rates Λ_*ij*_; since the log of these rates are parameterised as a log linear function of the predictors, we can obtain these rate through the coefficient estimates from the analysis and and predictors extended to include these additional locations. Specifically, we use this to predict introductions in regions in Guinea, for which no cases were reported (*n*=7) and for regions in neighbouring countries along the borders with Guinea or Liberia that remained disease free (*n*=18). To calculate the expected number of transitions from a particular phylogeographic location to any destination location, we sum over all destination locations (with and without cases, *n*=81). To obtain such estimates under different predictors or predictor combinations, we perform a specific analysis under the GLM model including only the relevant predictors or predictor combinations without the two-way random effects. For computational expedience, we performed these analyses, as well as the time-inhomogeneous analyses below, by conditioning on a set of 1000 trees from the posterior distribution of the main phylogenetic analysis (Lemey et al., 2014). We summarise mean posterior estimates for the transition expectations based on the samples obtained by our MCMC analysis; we note that also the value of *c* is sample-specific.

To consider time-inhomogeneity in the spatial diffusion process, we start by borrowing epoch modelling concepts from Bielejec et al. (2014). The epoch GLM parameterizes the instantaneous movement rate Λ_*ijt*_ from state *i* to state *j* within epoch *t* as a log-linear function of *P* epoch-specific predictors **X**_*ijt*_=(*x_ijt_*_1_*, …, x_ijtP_*)′ with constant-through-time, unknown coefficients ***β***. We generalize this model to incorporate time-varying contribution of the predictors through time-varying coefficients ***β***(*t*) using a series of change-point processes. Specifically, the time-varying epoch GLM models

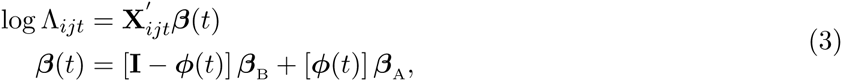
(3)
where ***β***_B_=(*β*_B1_*, …, β*_B*P*_)*t* are the unknown coefficients before the change-points, ***β***_A_=(*β*_A_1*, …, β*_A_*P*)*t* are the unknown coefficients after the change-points, diagonal matrix ***φ***(*t*) has entries (1_*t>t*_1 (*t*)*, …,* 1_*t>t*__*P*_ (*t*)), 1(*·*)(*t*) is the indicator function and **T**=(*t*_1_*, …, t_P_*) are the unknown change-point times. In this general form, the contribution of predictor *p* before its change-point time *t_p_* is *β*B*p* and its contribution after is *β*A*p* for *p*=1*, …, P*. Fixing *t_p_* to be less than the time of the first epoch or greater than the time of the last epoch results in a time-invariant coefficient for that predictor.

Similar to the constant-through-time GLM, we specify that *a priori* all *β*_B*p*_ and *β*_A*p*_ are independent and normally distributed with mean 0 and a relatively large variance of 4. Under the prior, each *t_p_* is equally likely to lie before any epoch.

We employ random-walk Metropolis transition kernels on ***β***_B_, ***β***_A_ and *T*.

In a first epoch GLM analysis, we keep the five predictors that are convincingly supported by the time-homogeneous analysis included in the model and estimate an independent change-point *t_p_* for their associated effect sizes: distance (*t_dis_*), within country effect (*t_wco_*), shared international border (*t_sib_*) and origin and destination population size (*t*_*pop*__*o*_ and *t*_*pop*__*d*_) change-points. To quantify the evidence in favour of each change-point, we calculate Bayes factor support based on the prior and posterior odds that *t*_*p*_ is less than the time of the first epoch or greater than the time of the last epoch. Because we find only very strong support for a change-point in the within country effect, we subsequently estimate the effect sizes before and after *t*_*wco*_, keeping the remaining four predictors homogeneous through time.

## Within-location generalized linear models

### Case counts

Ebola virus disease (EVD) case numbers are reported by the WHO for every country division (region) at the appropriate administrative level, split by epidemiological week. For every region and for each epidemiological week four numbers are reported: new cases in the patient and situation report databases as well as whether the new cases are confirmed or probable. At the height of the epidemic many cases went unconfirmed, even though they were likely to have been genuine EVD. As such, we treat probable EVD cases in WHO reports as confirmed and combine them with lab-confirmed EVD case numbers. Following this we take the higher combined case number of situation report and patient databases. The latest situation report in our data goes up to the epidemiological week spanning 8 to 14 February 2016, with all case numbers being downloaded on 22 February 2016. There are apparent discrepancies between cumulative case numbers reported for each country over the entire epidemic and case numbers reported per administrative division over time, such that our estimate for the final size of the epidemic, based on case numbers over time reported by the WHO, is on the order of 22 000 confirmed and suspected cases of EVD compared to the official estimate of around 28 000 cases across the entire epidemic. This likely arose because case numbers are easier to track at the country level, but become more difficult to narrow down to administrative subdivision level, especially over time (only 86% of the genome sequence have known location of infection).

We studied the association between disease case counts using generalized linear models in a very similar fashion to the framework presented above. A list of the location-level predictors we used for these analyses can be found in Table S2. We also employed SSVS as described above, in order to compute Bayes factors (BF) for each predictor. In keeping with the genetic GLM analyses, we also set the prior inclusion probabilities such that there was a 50% probability of no predictors being included.

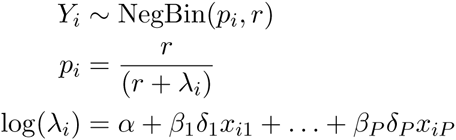

where *r* is the over-dispersion parameter, *δ_i_* are the indicators as before. Prior distributions on model parameters for these analyses were the same as those used for the genetic analyses whenever possible. We then employed this model to predict how many cases the locations which reported zero EVD cases would have gathered, that is, the potential size of the epidemic in each location.

### Computational details

To fit the models described above we took advantage of the routines already built in BEAST (https://github.com/beast-dev/beast-mcmc) but in a non-phylogenetic setting. Once again, posterior distributions for the parameters were explored using Markov chain Monte Carlo (MCMC). We ran each chain for 50 million iterations and discarded at least 10% of the samples as burn-in. Convergence was checked by visual inspection of the chains and checking that all parameters had effective sample sizes (ESS) greater than 200. We ran multiple chains to ensure results were consistent.

To make predictions, we used 50,000 Monte Carlo samples from the posterior distribution of coefficients and the overdispersion parameter (*r*) to simulate case counts for all locations with zero recorded EVD cases.

## Acknowledgments

The research leading to these results has received funding from the European Union Seventh Framework Programme [FP7/2007-2013] under Grant Agreement No 278433-PREDEMICS — Philippe Lemey and Andrew Rambaut, and ERC Grant Agreement No 260864 — Philippe Lemey, Andrew Rambaut and Marc A. Suchard. This work was supported by the European Unions Horizon 2020 research and innovation program (grant agreement no. 666100; EVIDENT) and the Directorate-General for International Cooperation and Development of the European Commission (service contract IFS/2011/272-372, EMLab) — Miles Carroll, David A. Matthews, Julian A. Hiscox, Antonino Di Caro, Roman Wlfel, Danny Asogun, Ekaete Alice Tobin, Joshua Quick, Nicholas J. Loman, Sophie Duraffour and Stephan Günther. European Unions Horizon 2020 research and innovation program (grant agreement No 643476.; COMPARE) — Marion Koopmans and Andrew Rambaut. National Institutes of Health (R01 AI107034, R01 HG006139 and R01 LM011827) and the National Science Foundation (IIS 1251151 and DMS 1264153) — Marc A. Suchard. National Health & Medical Research Council (Australia) — Edward C. Holmes. NIH AI081982, AI082119, AI082805 AI088843, AI104216, AI104621, AI115754, HSN272200900049C, and HHSN272201400048C — Robert F. Garry. The work in Liberia was funded by the Defense Threat Reduction Agency, the Global Emerging Infections System, and the Targeted Acquisition of Reference Materials Augmenting Capabilities (TARMAC) Initiative agencies from the U.S. Department of Defense — Gustavo Palacios. Bill and Melinda Gates Foundation (OPP1106427, 1032350, OPP1134076), Wellcome Trust Sustaining Health Grant (106866/Z/15/Z), Clinton Health Access Initiative — Andrew J. Tatem. This work was supported by the National Institute for Health Research Health Protection Research Unit in Emerging and Zoonotic Infections — Julian A. Hiscox. Key Research and Development Program (grant no. 2016YFC1200800) from the Ministry of Science and Technology of China — Di Lui. National Natural Science Foundation of China (NSFC, grant Nos. 81590760 and 81321063) — George F. Gao.

Colour-blind-friendly colour palettes by Cynthia Brewer, Pennsylvania State University (http://colorbrewer2.org). We gratefully acknowledge support from NVIDIA Corporation with the donation of parallel computing resources used for this research. Finally, we would like to recognize the contributions made by our colleagues who tragically died from Ebola virus disease whilst fighting the epidemic. In particular, we honor the memory of Dr. Sheik Humarr Khan and Nurse Mbalu Fonnie, whose careers were dedicated to viral hemorrhagic fever research.

## Supplementary Information

**Table S1.**
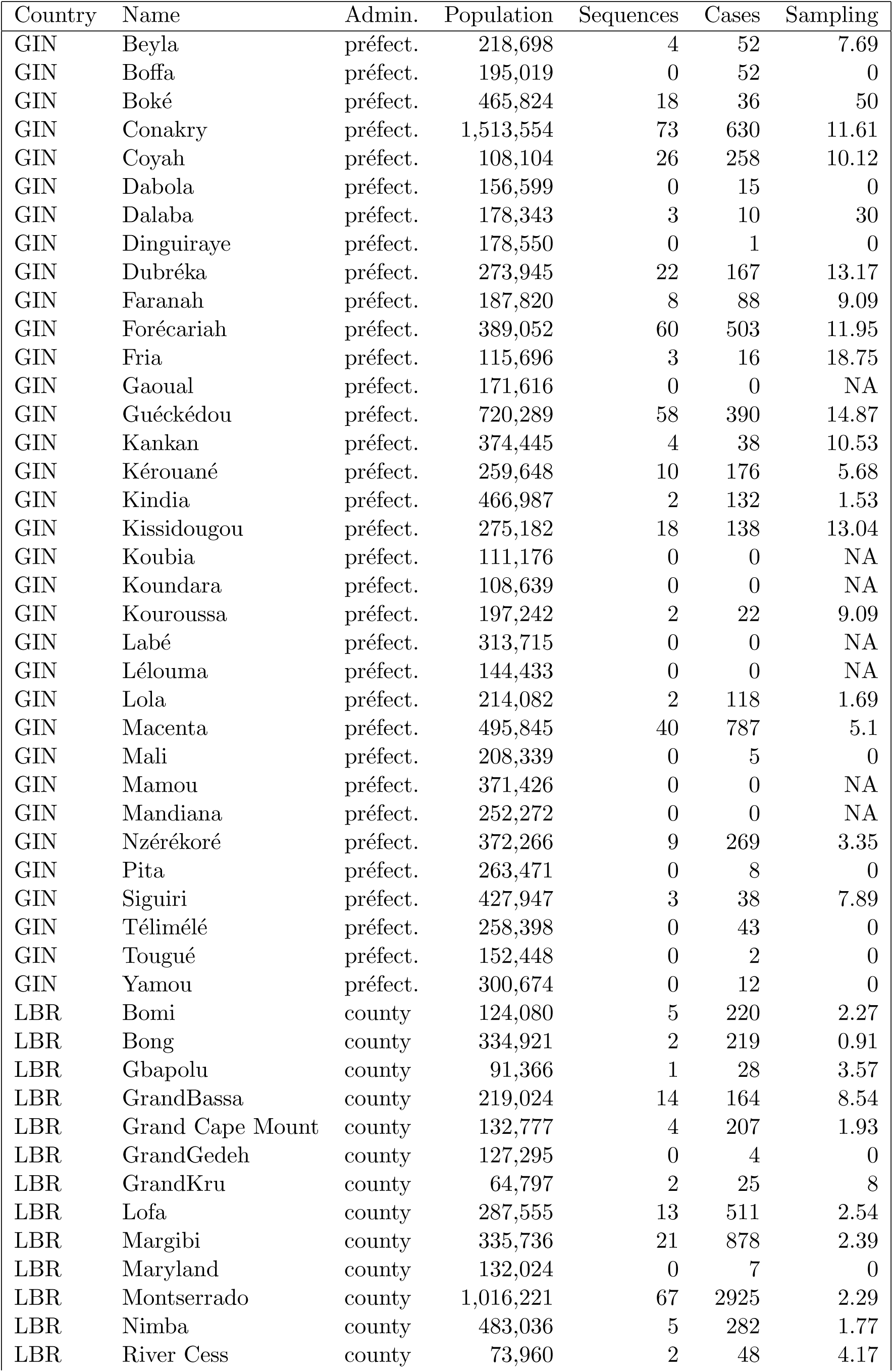
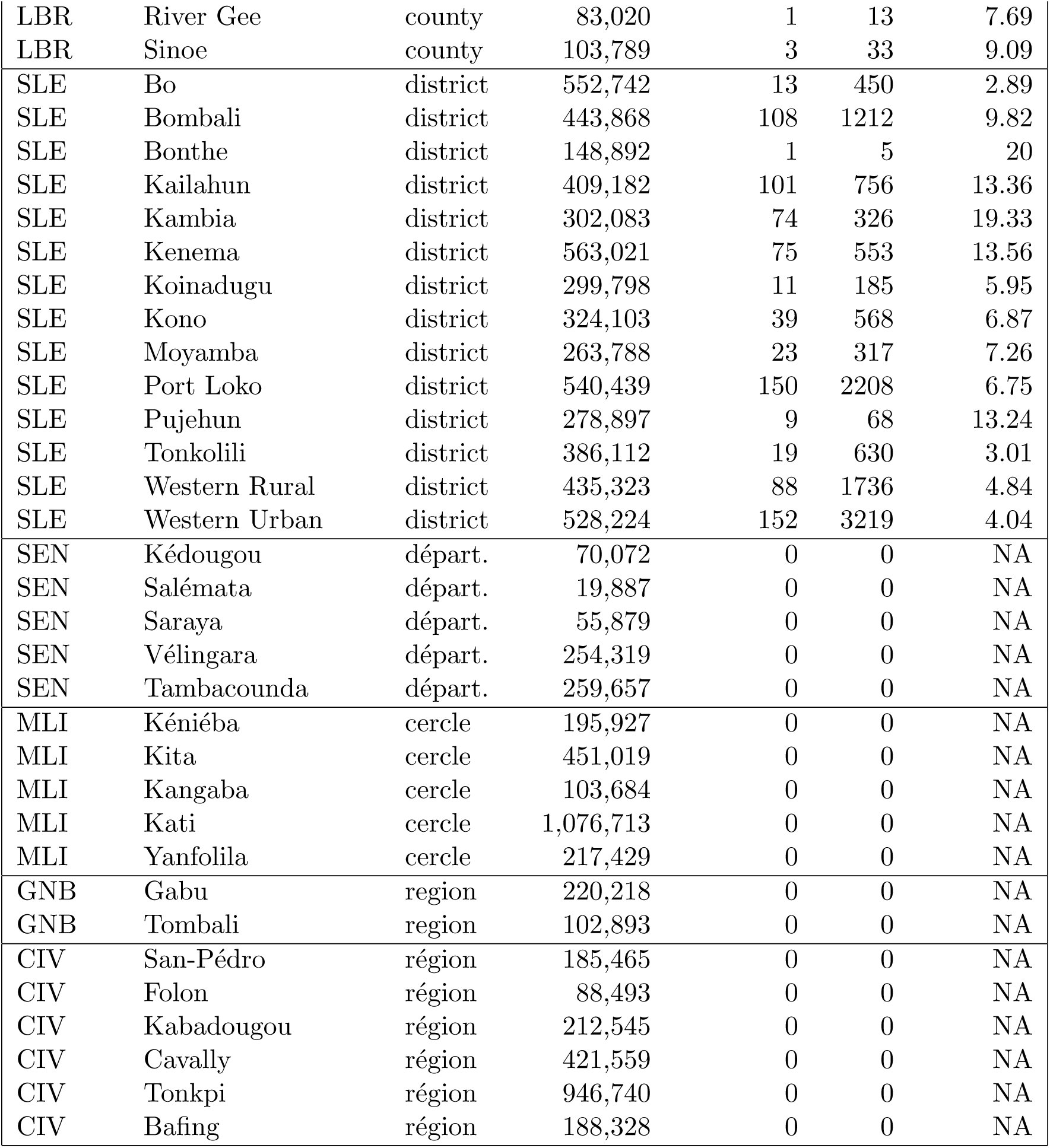
Number of cases and sampled sequences per region and country, where ‘Admin’ is the name of the administrative level used (‘préfect..’ being préfecture and ‘départ.’ being département) and ‘Sampling’ is sequences/cases × 100.

**Table S2.** Predictors included in the time-homogenous GLM.

| Predictor type | Abbreviation | Predictor description |
| --- | --- | --- |
| Geographic | Distances | Great circle distances between the locations' population centroids, log-transformed, standardized |
| Administrative | National | Two locations are in the same country |
| Administrative | IntBoSh | Location pairs that are in different countries and share a border |
| Administrative | NatBoSh | Location pairs that are in the same country and share a border |
| Administrative | LibGinAsym | Between Liberia-Guinea asymmetry |
| Administrative | LibSLeAsym | Between Liberia-Sierra Leone asymmetry |
| Administrative | GinSLeAsym | Between Guinea-Sierra Leone asymmetry |
| Demographic | OrPop | Origin population size, log-transformed, standardized |
| Demographic | DestPop | Destination population size, log-transformed, standardized |
| Demographic | OrPopDens | Origin population density, log-transformed, standardized |
| Demographic | DestPopDens | Destination population density, log-transformed, standardized |
| Demographic | orTT100k | Estimated mean travel time in minutes to reach the nearest major settlement of at least 100,000 people at origin, log-transformed, standardized |
| Demographic | destinationTT100k | estimated mean travel time in minutes to reach the nearest major settlement of at least 100,000 people at destination, log-transformed, standardized |
| Demographic | OrGrEcon | Origin Gridded economic output, log-transformed, standardized |
| Demographic | DestGrEcon | Destination Gridded economic output, log-transformed, standardized |
| Cultural | IntLangShared | Location pairs that are in different countries and share at least one of 17 vernacular languages |
| Cultural | NatLangShared | Location pairs that are in the same country and share at least one of 17 vernacular languages |
| Climatic | OrTemp | Temperature annual mean at origin, log-transformed, standardized |
| Climatic | DestTemp | Temperature annual mean at destination, log-transformed, standardized |
| Climatic | OrTempSS | Index of temperature seasonality at origin, log-transformed, standardized |
| Climatic | DestTempSS | Index of temperature seasonality at destination, log-transformed, standardized |
| Climatic | OrPrecip | Precipitation annual mean at origin, log-transformed, standardized |
| Climatic | DestPrecip | Precipitation annual mean at destination, log-transformed, standardized |
| Climatic | OrPrecipSS | Index of precipitation seasonality at origin, log-transformed, standardized |
| Climatic | DestPrecipSS | Index of precipitation seasonality at destination, log-transformed, standardized |

**Figure S1.**
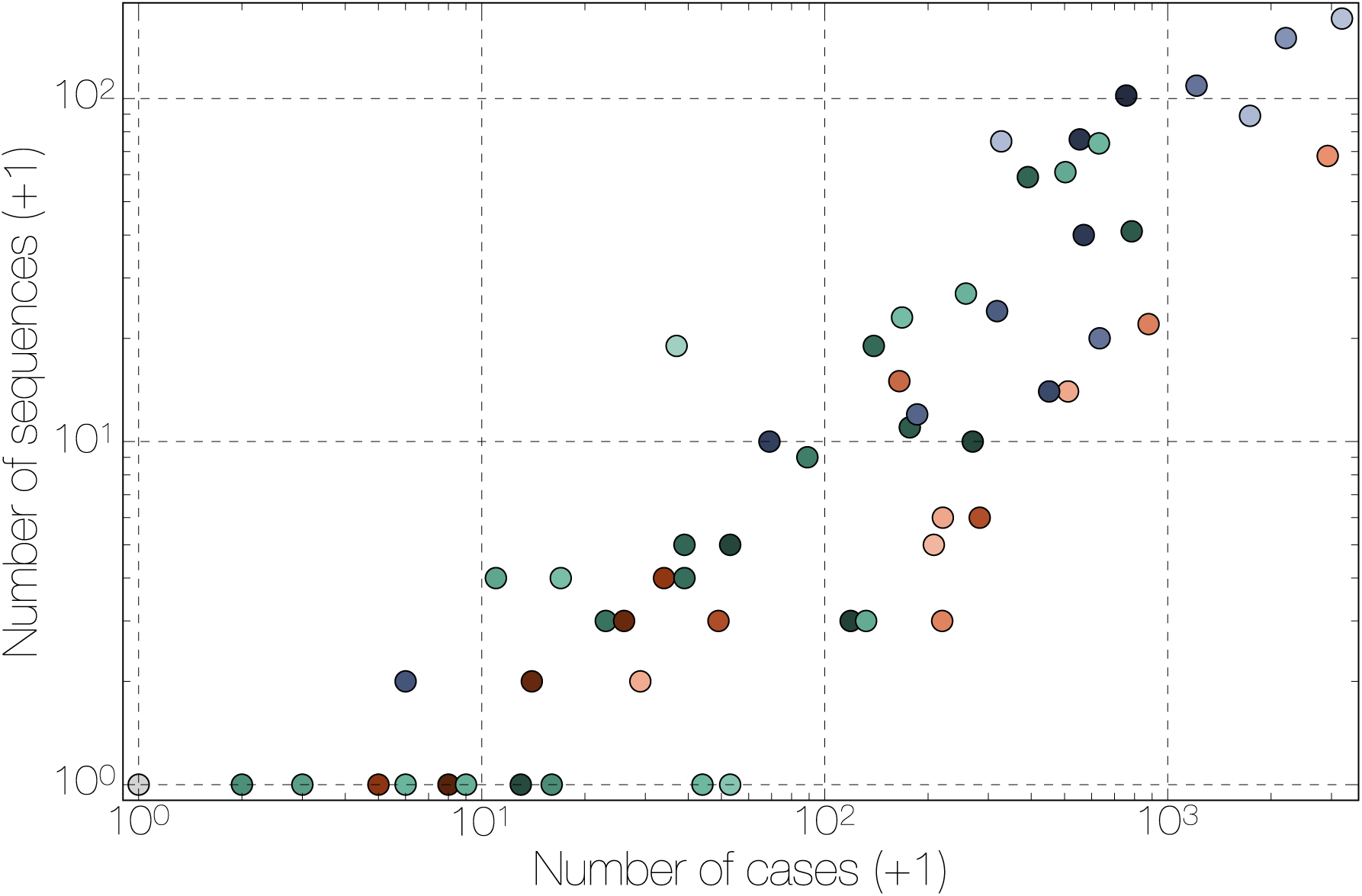
Correlation between number of cases and number of sequences for each location. A plot of number of EBOV genomes sampled against the known and suspected cumulative EVD case numbers. Regions in Guinea are denoted in green, Sierra Leone in blue and Liberia in red. Spearman correlation coe cient: 0.93.

**Figure S2.**
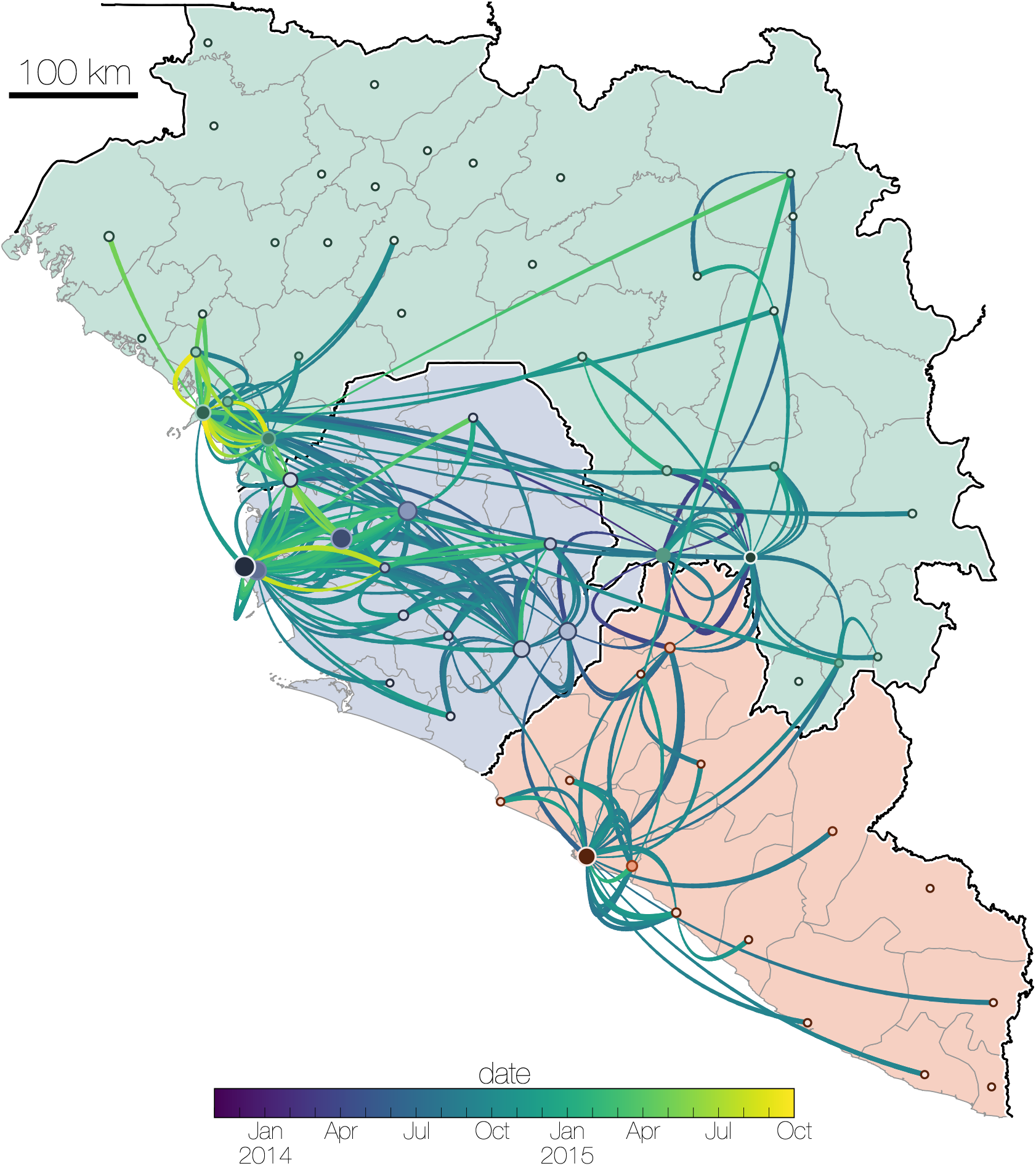
Dispersal of virus lineages over time. Virus dispersal between administrative regions estimated under the GLM phylogeography model (see Supplementary Methods). The arcs are between population centroids of each region, show directionality from thin end to thick end and are coloured in a scale denoting time from December 2013 in blue to October 2015 in yellow. Countries are coloured with Liberia in red, Guinea in green and Sierra Leone in blue.

**Figure S3.**
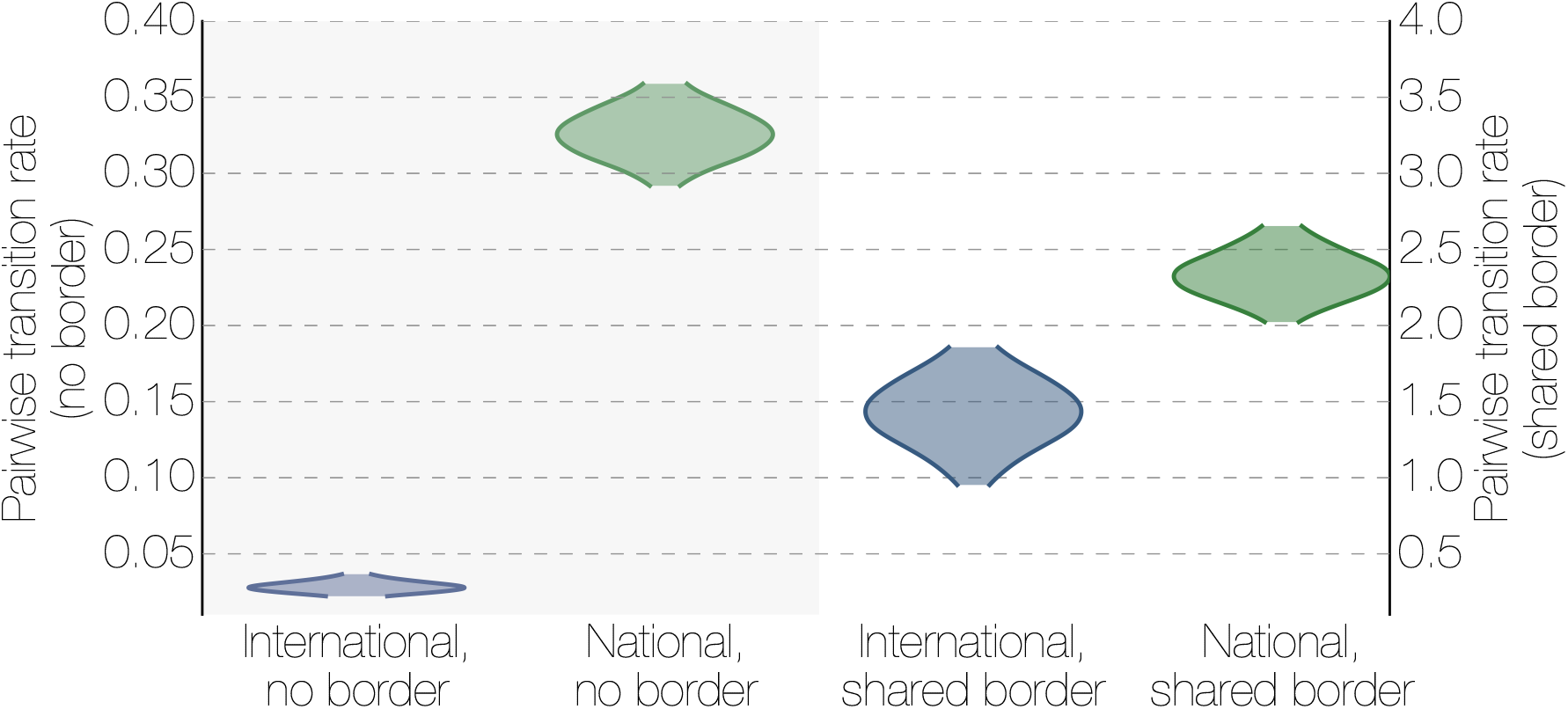
The effect of borders on EBOV migration rates between regions. Posterior densities of the migration rates between locations that share a geographical border (left) and those that do not (right) for international migrations and national migrations. Where two regions share a border, national migrations are only marginally more frequent than international migrations showing that both types of borders are porous to short local movement. Where the two regions are not adjacent, international migrations are much rarer than national migrations.

**Figure S4.**
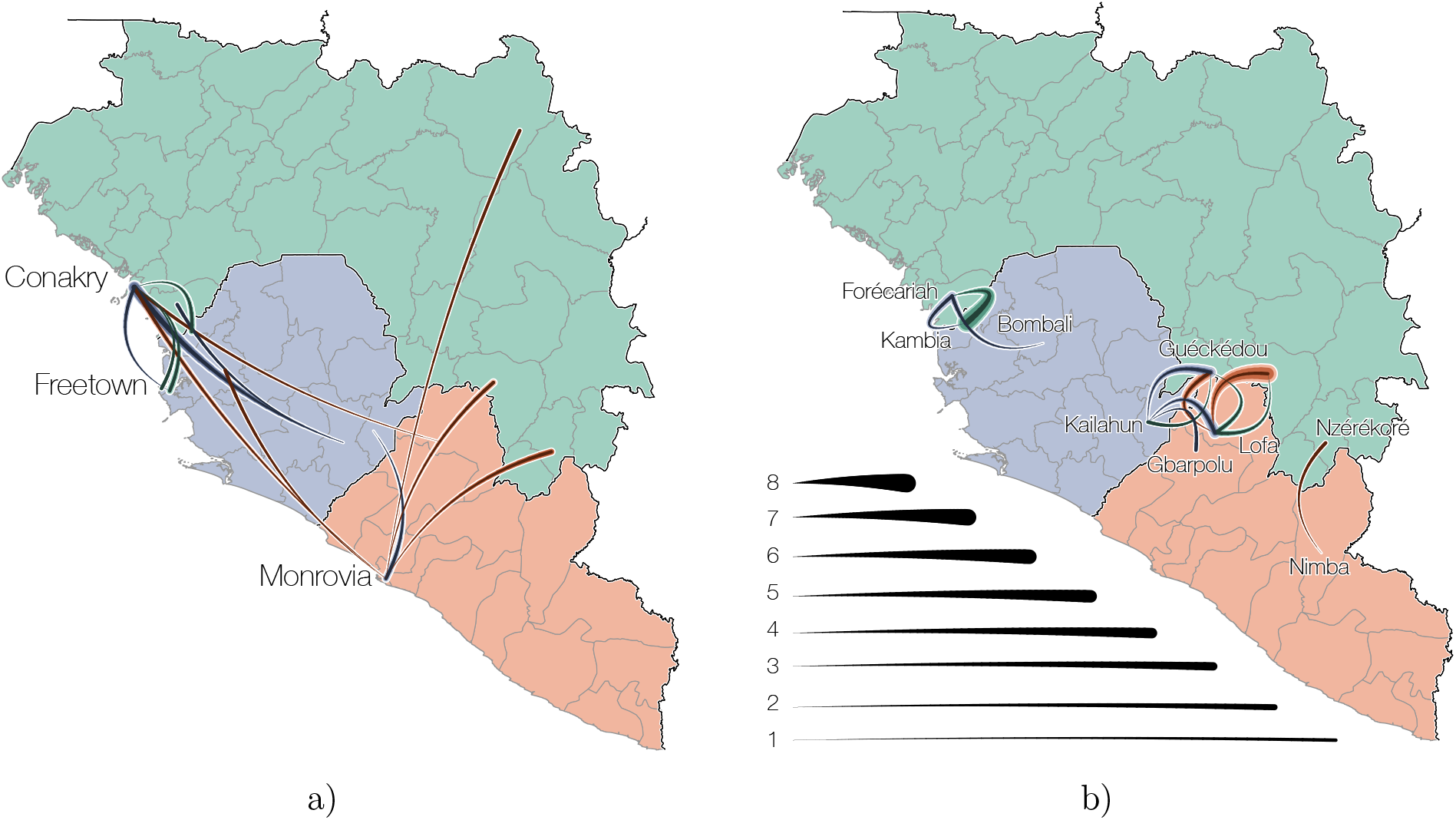
Summarized epidemic international migration history. All viral movement events between counties (Guinea, green; Sierra Leone, blue; Liberia, red) are shown split by whether they are between a) geographically distant regions or b) regions that share the international border. Curved lines indicate median (intermediate colour intensity), and 95% highest posterior density intervals (lightest and darkest colour intensities) for the number of migrations that are inferred to have taken place between countries.

**Figure S5.**
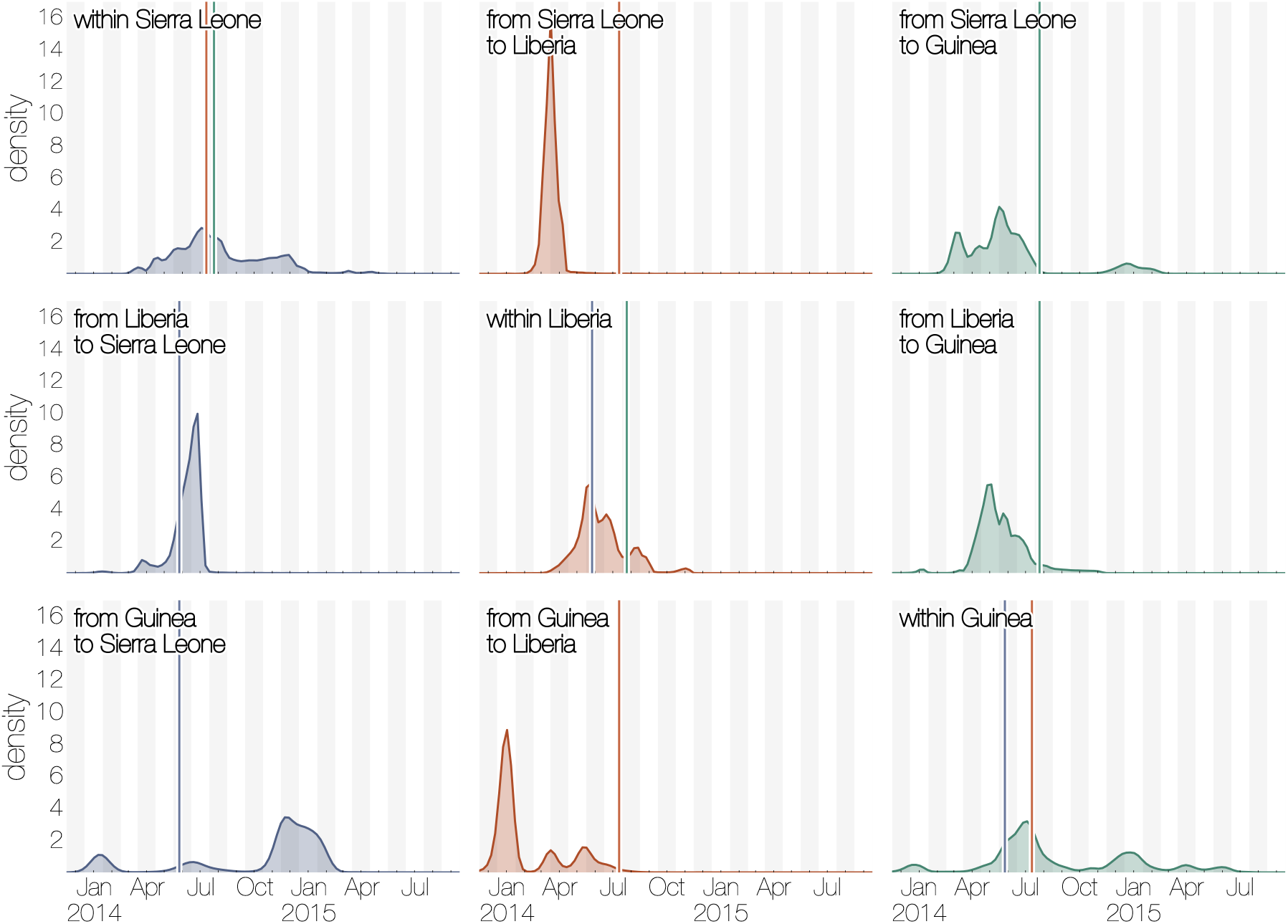
Summary of migration intensity over time in the region. Each cell shows the posterior probability density of temporal migration intensity. Vertical lines within each cell indicate the dates of declared border closures by each of the three countries: 11 June 2014 in Sierra Leone (blue), 27 July 2014 in Liberia (red), and 09 August 2014 in Guinea (green). Densities are rescaled and directly comparable across cells.

**Figure S6.**
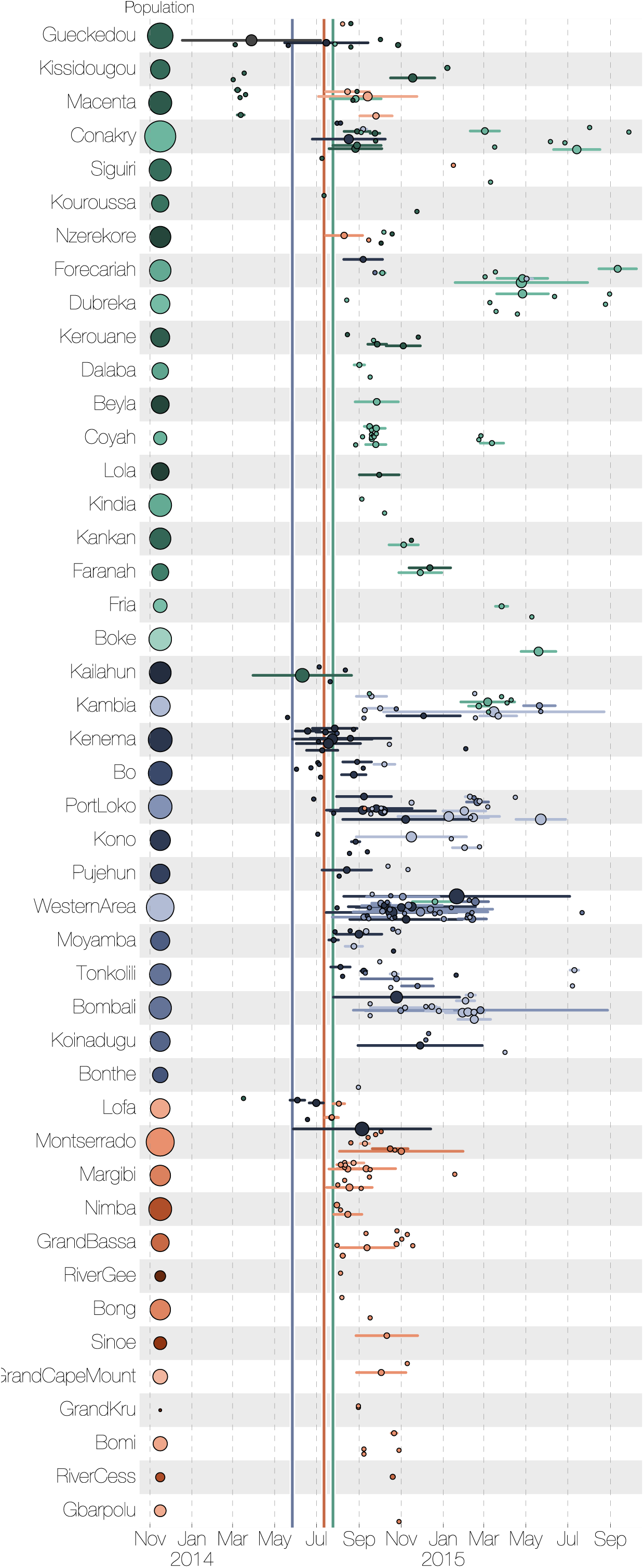
Region specific introductions, cluster sizes and persistence. Independent introductions into each administrative region and the size of each resulting cluster. The horizontal lines represent the persistence of each cluster from the time of introduction to the last sampled case. The areas of the circles in the middle of the lines are proportional to the number of sampled cases in the cluster. The areas of the circles next to the labels represent the population sizes of each administrative region. Vertical lines within each cell indicate the dates of declared border closures by each of the three countries: 11 June 2014 in Sierra Leone (blue), 27 July 2014 in Liberia (red), and 09 August 2014 in Guinea (green).

**Figure S7.**
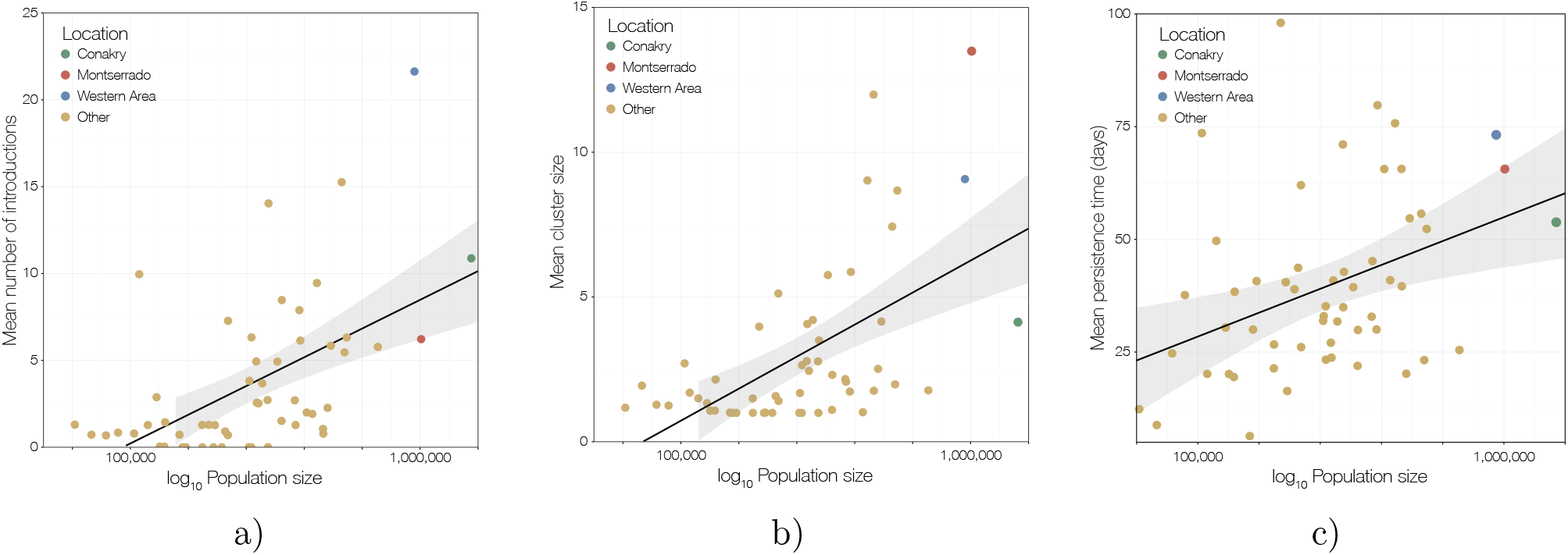
Relationship of cluster size, introductions and persistence to population size. a) The mean number of introductions into each location against (log) population sizes. The WesternArea (in Sierra Leone) received the most introductions, whilst Conakry and Montserrado were closer to the average. The association between population sizes and number of introductions was not very strong (*R*^2^=0.28, pearson correlation=0.54, Spearman correlation=0.57). b) The mean cluster size for each location plotted against (log) population sizes. The association here is weaker (*R*^2^=0.11, pearson correlation=0.35, Spearman correlation=0.57). c) The mean persistence times (per cluster, in days) against population sizes. A similarly weak association is observed (*R*^2^=0.12, pearson correlation=0.37, Spearman correlation=0.36). All computations based on a sample of 10, 000 trees from the posterior distribution.

